# *Bombus terrestris* Complementary Sex Determiner (BtCSD) is identified as a conserved hymenopteran sex determination region: evolution, breeding, and conservation implications

**DOI:** 10.1101/2025.08.12.669960

**Authors:** Kelley Leung, Frank Becker, Peter Šima, Roland Kreskóci, Elzemiek Geuverink, Thomas V. M. Groot, Bart A. Pannebakker, Joost van den Heuvel

**Author notes:** Author emails: Kelley Leung Frank Becker Peter Šima Roland Kreskóci Elzemiek Geuverink Thomas V. M. Groot Bart A. Pannebakker. (corresponding author) Joost van den Heuvel.

## Abstract

All Hymenoptera (bees, sawflies, ants, wasps) are haplodiploid. Haploid males develop from unfertilized eggs and diploid females from fertilized eggs. Many species also have single locus complementary sex determination (sl-CSD): hemizygosity of a CSD gene results in normal males and heterozygosity in normal females, but homozygosity typically yields sterile diploid males. Populations that lose CSD allelic diversity to critical level, increasingly produce detrimental diploid males. Emblematic of this problem are the bumblebees. Bumblebees are one of the fastest declining animal groups worldwide, with CSD diversity loss playing a likely role, especially in isolated populations. Presently utilized and potential future species in agriculture are also challenged by diploid males in production. Population management of both requires exact knowledge of the CSD locus. Here we report the identity of *Bombus terrestris* CSD, a bumblebee model that is also a major commercial pollinator. As CSD must be reciprocally homozygous in diploid males and heterozygous in diploid females, we whole-genome-sequenced these for four colonies. From this, we identified *B. terrestris* CSD (BtCSD) on Chromosome 2: a 26 Kb region with a long non-coding RNA region (lncRNA) and a proximate hypervariable region. Comparative analyses indicated conservation of this region across infraorder Aculeata, including other bumblebees, mason bees, and ants. This supports a deep-rooted ancestral origin for BtCSD despite a lack of homology with con-familial honey bee *csd* and rapid evolutionary divergence of insect sex determination cascades. Our analyses do not support the lncRNA region as functional CSD, as proposed for other taxa, because of inadequate variation to constitute distinct alleles. Rather, a downstream region is implicated for its massive structural variation and candidate coding genes including *Creld* and two copies of *aCOP*. We discuss how this identification of BtCSD has critical implications for hymenopteran evolution and bumblebee population management.

## Introduction

There are more than 150,000 described species in Hymenoptera (sawflies, ants, wasps and bees), with total diversity possibly exceeding a million species (Forbes et al., 2018). They all have haplodiploid sex determination. Haploid males arise from unfertilized eggs, and diploid females from fertilized eggs. Furthermore, numerous taxa including all Aculeata (bees, ants, stinging wasps) have complementary sex determination (CSD) (Whiting, 1943; Asplen et al., 2009; Leung & van der Meulen, 2022). In CSD, hemizygosity of any allele results in the development of functional males. Heterozygosity results in the development of functional females. Homozygosity however results in the development of aberrant diploid males that are typically sterile (Heimpel & de Boer, 2008). Hence any loss of CSD allelic diversity (e.g. through bottlenecks typical to isolated populations) increases diploid male production. These diploid males typically cannot produce functional female offspring, so a population is predicted to become smaller and more male-biased each generation, even possibly going extinct (diploid male vortex, or DMV) (Zayed & Packer, 2005; Leung & van der Meulen, 2022).

Measurement and management of CSD parameters to mitigate diploid male production is impossible without exact CSD loci to elucidate taxon-specific effects. Identifying CSD loci however has proven difficult across Hymenoptera, with candidates known in only a few species (ants *Vollenhovia emeryi* (Miyakawa & Mikheyev, 2015), *Linepithema humile* (Pan et al., 2024), *Ooceraea biroi* (Lacy et al., 2025); and orchard bee *Osmia bicornis* (Rönneburg et al., 2025)). A specific *csd* gene and its mechanism is only identified for *A. mellifera* (honey bee) (Beye et al., 2003; Hasselmann et al., 2008; Otte et al., 2023; Suzuki et al., 2023). Complicating matters further, species-specific characteristics can obscure under what circumstances the DMV might occur, e.g. mitigative factors like multiple mating and failures of diploid male development and reproduction (*A. mellifera* has both; Ratnieks, 1990) versus exacerbative factors like monogamous mating and functional diploid males (Leung & van der Meulen, 2022). Therefore, while often cited as a major risk to population survival in insect breeding and conservation, the mechanistic validity of a DMV is unknown for any CSD species.

A group uniquely positioned for resolving gaps of CSD knowledge, including roles in population health, are the bumblebees. Like all bees, the ~250 species of widely distributed bumblebees (*Bombus* spp.) are prescribed to be sl-CSD as well. (Duchateau et al., 1994; Leung & van der Meulen, 2022). Bumblebees are undergoing faster decline than other bees (Goulson et al., 2008; Williams & Osborne, 2009; Cameron & Sadd, 2020) (e.g. 3 times faster species-richness than other bees in the Netherlands (Van Dooren, 2019). The contribution of taxon-specific aspects of its sl-CSD mechanism is unclear. While there have been suggestions to use diploid males are population health indicator, it has not been incorporated in broader practice for bumblebee management (Whitehorn et al., 2009). For instance, a conservation model for bumblebees incorporates habitat loss, pesticides, and pathogens, but not genetic variation nor sex determination (Becher et al., 2018). However, bumblebees are not known to have compensatory mechanisms for diploid males. Queens are usually monogamous (Goulson, 2010). Unlike some other CSD species, *Bombus* queens also do not mate-discriminate against more closely related haploid males that would produce sterile diploid males (Bogo et al., 2018), nor against the diploid males themselves (Lecocq et al., 2017). Matings with diploid males very rarely result in any viable female offspring (Ayabe et al., 2004), and workers do not kill diploid male larvae or diploid-male producing queens, as in some bee species (Woyke, 1963, 1980; Duchateau et al., 1994; Alves et al., 2011; Vollet-Neto et al., 2017). Bumblebees therefore represent fewer complicating variables for direct measurement of CSD diversity impacts on population health, once a specific locus is known.

Meanwhile, a few bumblebee species are the most heavily produced group of agricultural pollinators after honey bees (Osterman et al., 2021). European buff-tailed bee *Bombus terrestris* has been imported worldwide; it and North American counterpart *B. impatiens* pollinate over 240 crops (IPBES, 2016). Diploid males are a known challenge in commercial breeding. Hives in which they occur are unusable because they displace pollinating female workers. This creates greater cost to breeders to replace these hives (Duchateau et al., 1994; Di Pietro et al., 2022). Such applied relevance means that the presence of sl-CSD in *B. terrestris* (Duchateau et al., 1994) It has also resulted in *B. terrestris*’s status as a model system with large, well-annotated genomics datasets, facilitating investigation for CSD locus identification.

Here, we report a single genomic region as *Bombus terrestris* CSD, BtCSD. CSD must be reciprocally homozygous in diploid males and heterozygous in diploid females. A single candidate matching these criteria was detected with de-novo whole genome sequencing and analyses of other available genomics datasets. Phylogenetic investigation indicated elements of remarkable evolutionary conservation across distantly related hymenopterans such as ants and *Osmia* mason bees, but not more closely related honey bees. In the discussion, we advise on applying knowledge of BtCSD towards future research on re-evaluating patterns of hymenopteran CSD evolution; conservation of wild bumblebee populations; managing present and future commercial populations; and further investigating *Bombus* CSD molecular mechanism(s).

## Methods

### Bumblebee material

A commercial producer supplied four, specially selected, early male-producing *B. terrestris* colonies. Typical outbred colonies do not produce males before the “switch point” (Duchateau & Velthuis, 1988), so all males present in these selected colonies were presumed diploid males (Di Pietro et al., 2022). Female workers, males, and queens were stored at 4°C in absolute ethanol until DNA extraction.

### Whole genome sequencing

Two hindlegs were collected from ten males and the queen of three *B. terrestris* colonies. Hindlegs of each individual were placed in separate tubes and the ethanol evaporated. They were then frozen in liquid nitrogen and mechanically ground using LabTIE Zirconia beads (Molgen, Veenendaal, The Netherlands). DNA was extracted using a standard cetyltrimethylammoniym bromide (CTAB, Sigma Life Sciences, St. Louis, MO, USA) method (https://openwetware.org/wiki/Maloof_Lab). DNA library preparation was done with Hackflex (Gaio et al., 2022). Briefly, DNA samples were quantified using SYBR green (Invitrogen, USA) and diluted to 0.2-0.25 ng/μL. 2 μL of diluted DNA was tagmented and adapter-ligated in a 3 μL reaction using an Illumina Nextera DNA Flex Library Prep Kit (Illumina, San Diego, CA, USA). Libraries were amplified using Illumina TruSeq primers and Robust 2G enzyme (Kapa Biosystems, Cape Town, South Africa). Libraries were size-selected for 300-500bp fragments using solid phase reversible immobilization beads (SPRI, SpeadBead Magnetic Carboxylate Modified Particles, Cytiva, Marlborough, MA, USA). Selected libraries were paired-end sequenced by Novogene Europe (Cambridge, United Kingdom), targeting an average depth of 15X whole genome coverage per diploid male, and 50X coverage per queen.

### Bioinformatic analyses

In addition to our sequenced libraries, we used data from the Short Read Archive to quantify variation in diploid workers (PRJEB52013 (Barribeau et al., 2022), PRJEB49221 (Kardum Hjort et al., 2022)) and haploid drones (PRJNA628944 (Colgan et al., 2022)) (Supplemental File 1). Raw reads were trimmed using Trimmomatic (v 0.33, adapters.fa:2:30:10 LEADING:3 TRAILING:3 SLIDINGWINDOW:4:15 MINLEN:50, using adapter sequences CTGTCTCTTATACACATCTGACGCTGCCGACGA and GTCTCGTGGGCTCGGAGATGTGTATAAGAGACAG (Bolger et al., 2014)). Trimmed reads were then mapped to the reference genome of *B. terrestris* (iyBomTerr1.2, (Crowley et al., 2023)) using bwa-mem2 (Li & Durbin, 2009). Both single and paired-end reads were mapped after trimming. Mapped reads filtered for quality and alignments sorted for position resulted in a sorted bam file using samtools (v. 0.1.20, -q 20, (Li et al., 2009)). Duplicates were removed with Picard tools (v 2.8.2, (Broad Institute, 2019)) and Freebayes-parallel was used to call variants (-F 0.25, (Garrison & Marth, 2012)). Repeats were removed using bedtools intersect (Quinlan & Hall, 2010).

### Population genetic parameters

We identified heterozygous variants with the worker datasets and false positive sites of heterozygosity with the drone dataset. We used the R library vcfR (Knaus & Grunwald, 2017) in R (4.0.2; (Team, 2024) to calculate minor allele frequencies (MAF), variant quality, and mean depth over all samples. Variants were kept if MAF>0.02, 10 < mean depth <50, variant QUAL >5 (Kardum Hjort et al., 2022) and MAF>0.02, 5 < mean depth <25, variant QUAL >5 (Barribeau et al., 2022). Heterozygosity for all variants was conservatively counted for the drone dataset without MAF of read depth cutoffs. We masked the heterozygosity in both datasets if variants were in 10 bps proximity of drone false heterozygous sites. A variant was called heterozygous if two alleles differed from each other, using the GT field from the function extract.gt(vcf) in the vcfR package. For the diploid males, we calculated pairwise genetic distance as the difference in genotype calls in windows of 100 adjacent variants that were called heterozygous in the queens, using genotype calls from the same GT field (3 < mean depth <13, variant QUAL >1). For the MDS, genetic distance was calculated using alternative reads frequencies. These frequencies were obtained from the read depths, where alternative read depth was taken as the ‘AO’ field from extract.gt(), and reference read depth was taken from the ‘RO’ field from extract.gt(). Once a distance matrix was produced, the MDS plot was produced with cmdscale (R base, 4.0.2;(Team, 2024).

### De novo assembly of drones

We used whole genome data from drones (Colgan et al., 2022) to de-novo assemble the CSD region (see Supplemental File 2 for code). First, sorted and deduplicated bam files were used to extract raw reads aligning around the putative CSD locus using “samtools view bams/$pop.sort.bam “NC_063270.1:16252000-16284000” | awk ‘{print $1}’ | sort | uniq”. Seqtk (https://github.com/lh3/seqtk) was used to extract a list of reads from raw fastq files. Reads were assembled using spades (v3.13.0, (Bankevich et al., 2012) We used blastn (Camacho et al., 2009) to identify scaffolds as the CSD locus and only kept those that showed a hit with NC_063270.1:16250000-16290000 as a reference. We then produced an assembly using these reads. We used this 40 Kb CSD region to search for reads in the raw data, using a kmer search (bbduk.sh, k=31, hd. https://sourceforge.net/projects/bbmap/.) Those hits were also used for spades assembly to produce an alternative assembly. These two assemblies were merged using quickmerge (Chakraborty et al., 2016) with the first assembly as ‘reference’ and the bbduk second assembly as query genome (labelled “merged1”). The resulting merged assembly was again filtered using blastn using the 40 Kb CSD region, but the output was filtered for 90% identity and hits of at least 1000 bp length. As many assemblies were still discontinuous, we isolated sequence on the ‘edges’ of the scaffolds by taking the first and last 300 bp of each scaffold (seqkit –r 1:300 and –r −300:−1) and used these sequences to search for matching raw data (bbduk.sh, same parameters above). These reads were assembled for a third assembly using spades. The resulting scaffolds were merged with the previous “merged1” assembly to produce a final merged assembly. From this final assembly, only sequences that contained both the upstream gene LOC125387186 and the downstream gene *Creld* (LOC100650350) were kept. In total this resulted in 17 assemblies from 51 samples (Supplemental File 4).

### Kmer synteny blocks

Kmer synteny blocks were identified using a custom R script (Supplemental File 3). Briefly, all kmers were identified in the 17 haploid assemblies and in the putative CSD region of the reference genome (Supplemental File 4). Only unique kmers were kept. Common kmers were identified between assemblies and only those which were found more than once (but not in excess in the number of assemblies) were kept, to remove repeats. Kmers were grouped into blocks if the distance between them was less than 20 basepairs. To each common kmer, sample-specific block numbers were added. Common kmers were grouped into numbered blocks, with numbers indicating proximity on the genome. Blocks with more than 400 common kmers were kept as synteny blocks. The relative position of these blocks (start to end) was then plotted.

### Long distance PCR

Using the 18 haploid assemblies, two primer pairs were developed using primer-blast: yF1 – GACCGTCTGCGTTGAAATCG, hr1 – GTGATAGCGCATGGGAGGAA and yF2 – AGCCTTCGGAAATCAGTCCG, hr2 – ATTTCCTCCATGTGCGGCTT. Pair 1 is positioned at 16,275,511-16,283,018 and pair 2 at 16,275,487-16,283,337 in the reference genome. PCR was performed using PCRBIO VeriFi™ Mix. The PCR mix was 12.5 µL Verifi Mix, 1 µL (10µM) forward primer, 1 µL (10µM) reverse primer, 1 µL DNA template and 9,5 µL MQ. The PCR cycle was 1 min of 95°C degrees denaturation and 35 cycles of 95°C denaturation (15s), 60°C degrees annealing (15s), and 72°C degrees extension (5.5 minutes). Ethidium bromide 1% gel electrophoresis was used to visualize PCR results. Following SPRI bead cleanup, PCR products were sent to Plasmidsaurus for long read sequencing using Oxford Nanopore Technology.

### Amplicon sequencing

The structural variation might have obscured or produced false heterozygosity sites in the alignment of the short reads. Therefore, we used amplicon sequencing of four conserved regions that contained variants in queens for Colonies 1 to 3 using 1.8 Kb – 2.5 Kb length products. We amplified these products for each queen, 10 workers and 12 diploid males of each family. We also did this analysis for the fourth family, which was not included in the whole genome sequencing. For amplicon sequencing, primers were developed using the reference genome (regions of low variation in the sequenced populations visualized in IGV (Robinson et al., 2011)). Four primer pairs were developed (Ampl1F: Ftail-ATCCTGTGGCTAGACTCGGT, Ampl2F: Ftail-GCCAATTTAGCACGCGAGAG, Ampl3F: Ftail-ATCACCTGTGCACCCATCAC, Ampl4F: Ftail-ATGATTAGGTGGCTTGGGGC, Ampl1R: Rtail-CCTCTTTAGCTCCCTGTGGC, Ampl2R: Rtail-CGAATTTCGCTGCACCTACG, Ampl3R: Rtail – TCACGAAATGCGACACGTCA, Ampl4R: Rtail –TGTTCAGCGTTTCAATCACG). Primers contained Ftail TTTCTGTTGGTGCTGATATTGC and Rtail ACTTGCCTGTCGCTCTATCTTC to enable library prep according to manufacturer protocol with the Ligation sequencing amplicon prep kit (V14, SQK-LSK1114) (Oxford Nanopore Technologies, Oxford, UK). Each of the 4 PCR reactions was 4 µL 5x GoTaq buffer, 0.8 µL DNTPs, 0.2 µL GoTaq polymerase (Promega, Madison, WI, USA), 0.8 µL (10µM) forward primer, 0.8 µL (10µM) reverse primer, 2 µL DNA template, and 11.6 µL MQ. The PCR cycle was 5 min of 95°C denaturation, and 33 cycles of 95°C denaturation (30s), 62°C degrees annealing (30s), and 72°C degrees extension (2.5 minutes). To start library prep with roughly equimolar PCR product, electrophoresis gels were inspected. Because bands were weaker for amplicons 3 and 4, we used 4 µL for amplicons 1 and 2, and 6 µL for amplicons 3 and 4. We then measured and then equalized DNA concentrations of all PCR product pools with PicoGreen (Thermo Fisher Scientific, Waltham, MA, USA). After barcode ligation, DNA concentrations were measured and normalized. The last end prep was sequenced on a Mk1C Nanopore device (Oxford Nanopore Technologies, Oxford, UK), which resulted in 965 Mb of data of fastq passed bases attributed to specific barcodes. The resulting fastq files were aligned to a partial reference genome (NC_063270.1:16,000,000 – 16,500,000) using minimap2 (v2.1, (Li, 2018). Samtools (v. 0.1.20, -q 20, (Li et al., 2009)) was used to sort reads and write out in bam format. Picard tools (v 2.8.2 (Broad Institute, 2019)) was used to add read groups. Variant calls were made with freebayes-parallel (Garrison & Marth, 2012). Variants were filtered using vcfR (Knaus & Grunwald, 2017). Only variants that were heterozygous in >5 individuals (0.4 < allele frequency < 0.6 was considered heterozygous), were biallelic, and of log variant quality >8 were kept. Only individuals with an average coverage over 200x for these variants were considered.

### Homology blast search

We used blast to investigate CSD loci region similarity among different hymenopteran taxa. We used sequences NC_063270.1: 16,210,000-16,290,000 (*B. terrestris*), NC_090131.1: 26,510,000-26,590,000 (*Linepithema humile*), NC_060216.1: 14,375,000-14,425,000 (*Osmia bicornis*) and NC_063270.1|:16210000-16290000 (*Ooceraea biroi*). We used blastn (Camacho et al., 2009) to blast all sequences to each other and identify regions larger than 40 basepairs (-word_size 10). A homology plot was made to identify these regions.

To detect the CSD loci region across Hymenoptera, we used peptide sequences of the genes flanking *B. terrestis* CSD (BtCSD) (THUMPD3 XP_003393947.1, CRELD XP_048267559.1 and aCOP XP_048267555.1) in tblastn against available high-quality genome assemblies. The selected databases (Table S1) included all major groups of the infraorder Aculeata as well as known sl-CSD species in the infraorder Proctotrupomorpha.

## Results

Whole genome sequencing identified variation within three colonies of one queen and 10 diploid males. For each queen there was a large degree of heterozygosity (423,527, 392,101 and 421,972 heterozygous variants for Colonies 1, 2 and 3 respectively). Calculated genetic distance (Supplementary Figure S1) indicated high relatedness between diploid males and their queens. The diploid males also had a high average number of heterozygous variants (109,578, 24,100 and 77,511 for each of the three colonies). This confirmed diploid (excluded haploid) status for the male samples. Lower realized sequencing coverage resulted in fewer heterozygous sites in diploid males relative to queens, and in queen 2 relative to other queens.

For all heterozygous variants in each queen, the pairwise genetic variation (π) between the 10 diploid males in each family identified BtCSD (Figure 1). That is, in diploid males, the CSD locus should be of low pairwise genetic variation relative to the rest of the genome. One such region was recovered for Colonies 1 and 3 on Chromosome 2, and two such regions were recovered for Colony 2 on Chromosomes 2 and 16.

**Figure 1.**
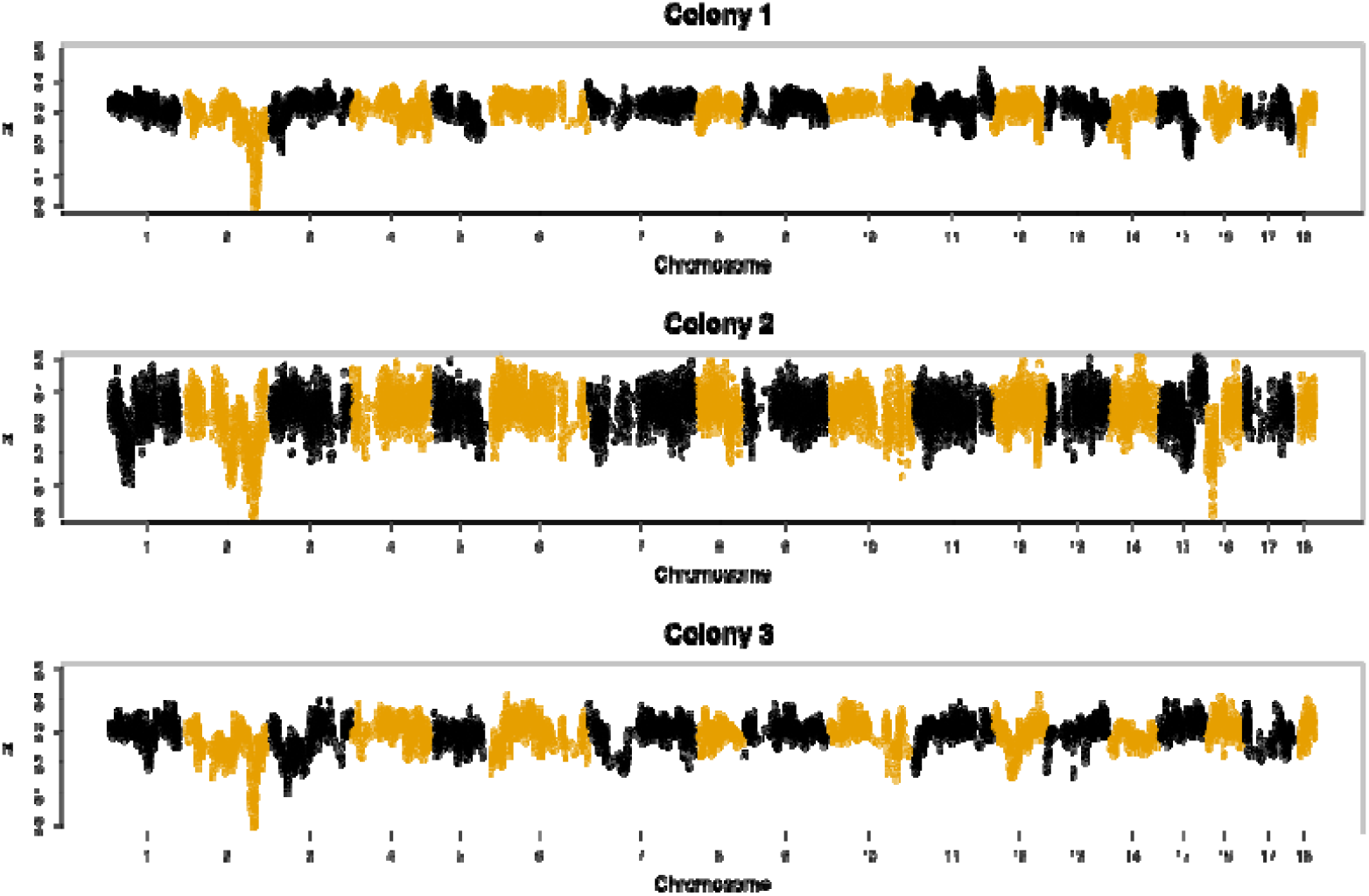
Genetic variation among 10 DMs within three colonies of *B. terrestris*. Y axis indicates mean pairwise genetic variation p. Numbers on the X axis indicate chromosome. Mean pairwise genetic variation is quantified for 100 consecutive positions that were heterozygous for the queen, with steps of 25 positions (75% overlap between neighboring windows).

Regions of average low heterozygosity in the diploid males were identified in each colony (heterozygosity<0.05, grey area Figure 2), and those overlapped between position 16,261,994 and 16,287,800 on Chromsosome 2 (green area Figure 2) for all colonies. Average heterozygosity 2.5 Kb windows for two studies that whole genome sequenced *B. terrestris* workers (Barribeau et al., 2022; Kardum Hjort et al., 2022) was highest between 16,278,500 and 16,281,500 on Chromsosome 2, further validating this region as BtCSD. In addition to this region’s high average heterozygosity, the number of variants is also highest there for both worker studies (Figure 2), as expected for a CSD locus. Furthermore, our identified BtCSD locus is homologous to the CSD locus previously reported in the ants *L. humile* (Pan et al., 2024) and *Ooceraea biroi* (Lacy et al., 2025) and in another bee, *Osmia bicornis* (Rönneburg et al., 2025). This corroborates this region as CSD in *B. terrestris* and indicates that a CSD locus is evolutionary conserved among bees and ants.

**Figure 2.**
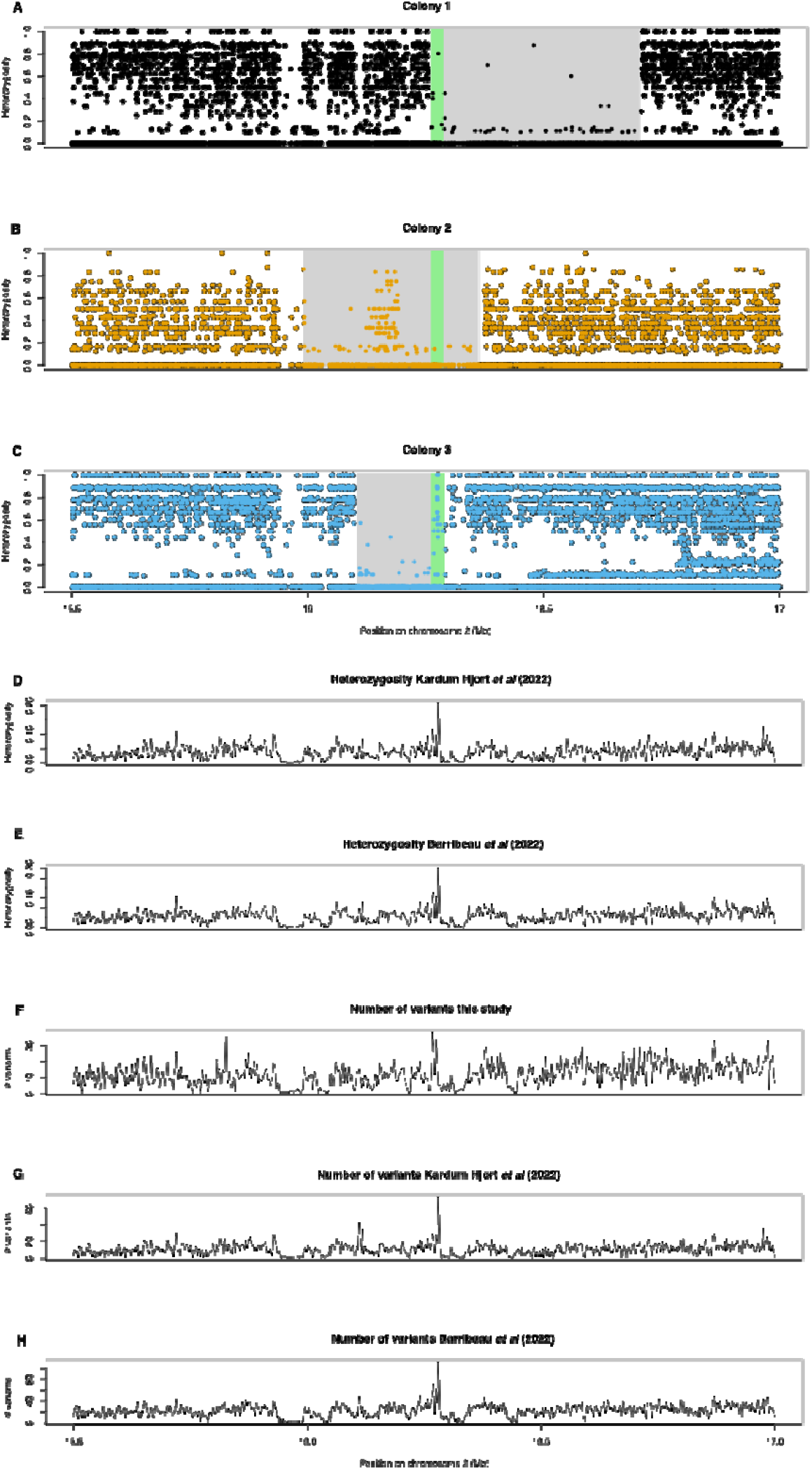
Genetic variation around the CSD locus. Panel A-C: heterozygosity in Colonies 1 to 3. Each dot represents the average heterozygosity per variant, the grey area indicates a region with average low heterozygosity among 50 adjacent variants. Panel D-E: heterozygosity in 2.5 Kb windows in workers, from (Kardum Hjort et al., 2022) and (Barribeau et al., 2022) who performed whole genome sequencing. Panel F-H: Number of variants in this study (F) and in (Kardum Hjort et al., 2022) (G) and (Barribeau et al., 2022) (H). The overlapping region of low heterozygosity in Colonies 1-3 overlap with a highly variable region in the two studies of (Kardum Hjort et al., 2022) and (Barribeau et al., 2022), both for heterozygosity per locus and the number of variants present within a 2.5 Kb window.

Although the average heterozygosity was low within the 26 Kb candidate region, Family 1 and Family 3 showed some individual loci with high heterozygosity. Visual inspection of these regions for the three colonies (Supplemental Figure S2) showed structural variation within BtCSD. This was indicated by loss of coverage in the alignments; a relatively high abundance of reads that only partially aligned; and reads with abnormal sequence length between left and right read per paired-end.

De-novo assembly on shorts reads of whole genome sequenced haploid males (Colgan et al., 2022) better reconstructed this variation. Comparison of synteny blocks in the 17 assembled CSD loci and the reference genome sequence (iyBomTerr1.2) (Figure 3) clearly indicated structural variation. For instance, individual SRR11640395 showed an inverted sequence of synteny blocks ‘rmly’. Some synteny blocks were found in only a few individuals, such as ‘woe’, ‘i’ or ‘g’. In our whole genome sequence alignment, missing coverage was potentially due to these insertions and deletions. Lastly, synteny blocks differed greatly in their relative distances (Figure 3).

**Figure 3.**
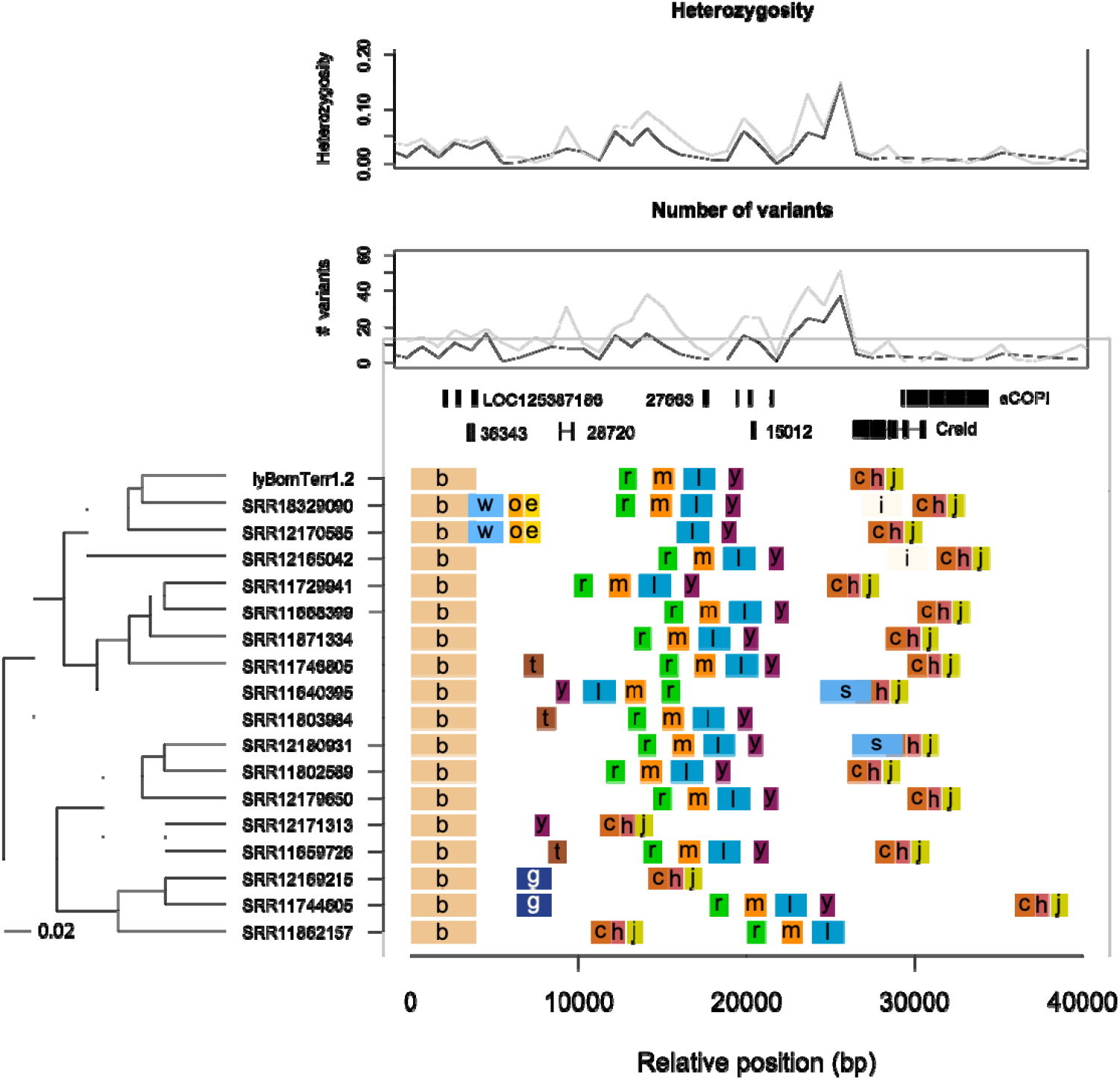
Structural variation at the CSD locus. Variation of the CSD locus of de-novo assembled sequences of 17 haploid males is shown (data taken from (Colgan et al., 2022)). A synteny block indicates a region that is highly similar between at least two individuals. The upstream and downstream regions are highly conserved. Gene models on top of the figure are indicated by gene or locus names, where a number is listed, it corresponds to an Ensembl gene number. Individuals are ordered along the Y axis according to their genetic distance, based on kmer dissimilarity (Jaccard dissimilarity of presence / absence of kmers), clustered using hierarchical clustering. Samples are named after their NCBI accession number. Heterozygosity and number of variants are indicated (per 1 Kb window) above the gene models. Variation peaks just before the Creld gene.

Because short reads were used, we needed to exclude this variation being due to incomplete (mis)assembly. Therefore, to validate the length differences between the synteny blocks, we performed long-distance PCR of synteny block ‘y’ to ‘h’ (Figure 3) on the queen and two diploid males from an additional Family 4. Surprisingly, only one relatively short band was recovered (~7 Kb in the queen, ~10.5 Kb in the two diploid males) (Supplemental Figure S3). Because the two diploid males were sons of the queen, the longer DNA molecule should have also been present in the queen. We hypothesized that the larger band was not visible using gel electrophoresis because the shorter product amplified more efficiently.

Long-read sequencing and alignment to identify similar regions between the two alleles found that, as expected, the first 800 and last 2000 base pairs aligned well, while the other sequences could not be aligned (Supplemental File S1). Both PCR products started in synteny block ‘y’ and ended in synteny block ‘h’, although the length of the product differed considerably. Sequencing recovered the long fragment in the queen sample as well, but at lower coverage than the shorter product. The long-distance PCR therefore corroborated large length differences in the BtCSD locus across individuals.

We then amplified four additional regions in 86 individuals within BtCSD to further validate differences in heterozygosity between diploid males and workers in four colonies. Similar to the short-read analysis, amplicon sequencing revealed that diploid males had very low heterozygosity levels, while queens and workers showed high levels of heterozygous sites for BtCSD (Figure 4). The positions of those heterozygous sites in diploid males were also similar, compared to the short-read analysis. For instance, in colony 1, heterozygous sites were found upstream (at LOC125387186) while for colony 3 heterozygous variants were identified downstream of BtCSD (at *Hcf*). The few heterozygous sites found in diploid males within the CSD region indicates that some variation exists between functional similar alleles. This analysis further corroborates with region as BtCSD, as workers show heterozygosity where diploid males lack it.

**Figure 4.**
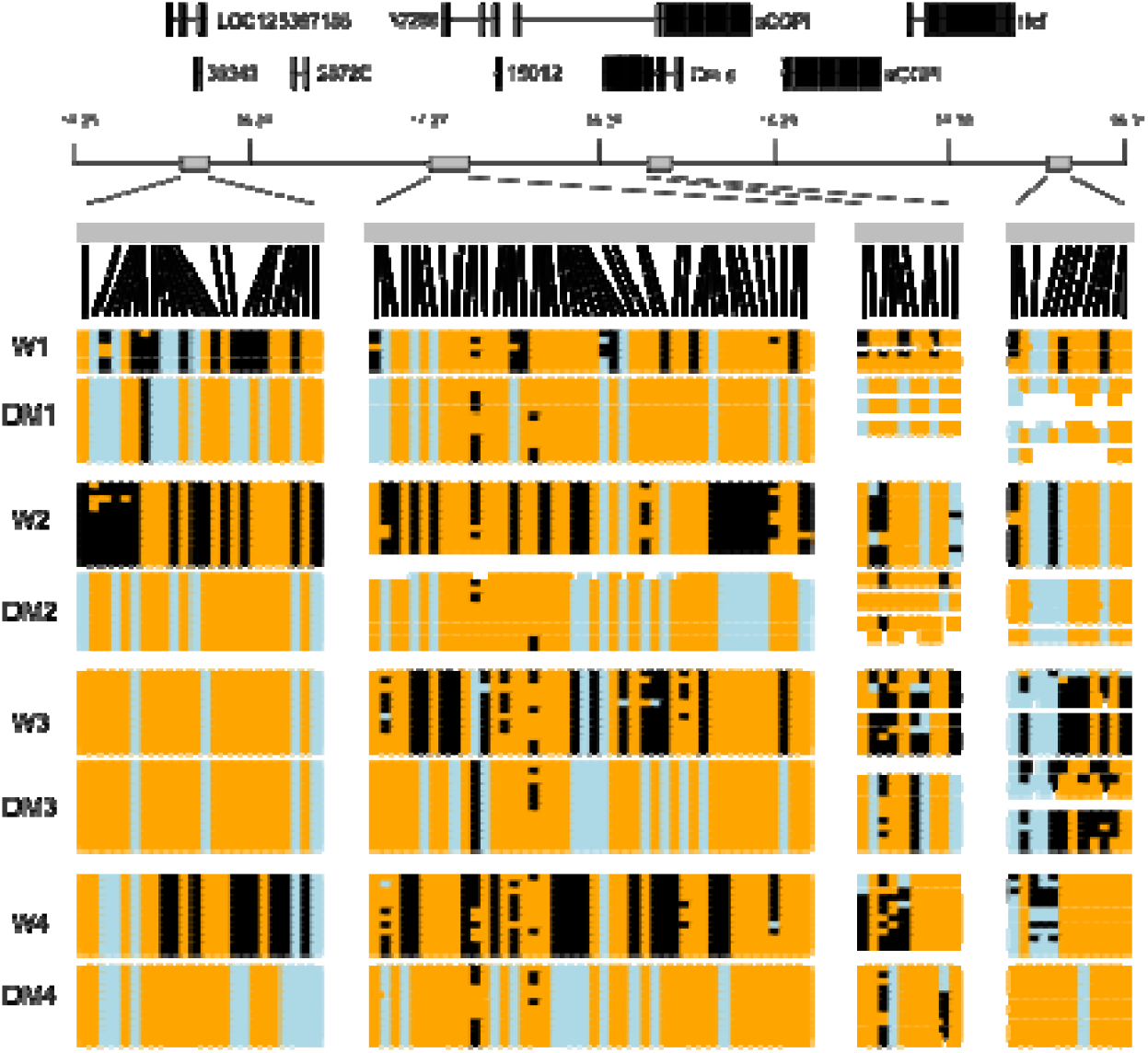
Amplicon sequencing results. From left to right 90 loci are indicated spread over 4 PCR products indicated by the grey boxes above. Green boxes indicate genes around the CSD locus, while numbers along that line indicate positions on Chromosome 2 in Mb. For each sample types (W=worker, DM = diploid male) we called genotypes as either homozygous reference (orange), heterozygous (black) or homozygous alternative (light blue). As expected, DMs show much lower degrees of homozygous genotypes, while workers are more likely heterozygous.

As the CSD candidates of *L. humile, Ooceraea biroi, Osmia bicornis* and *B. terrestris* are in the same genomic region, it was of interest to identify homologous genetic elements among these species. Blasting the four regions against each other notably showed that the larger structural variant in *B. terrestris* lacked any similarity with these species. The relative distance between the upstream and downstream elements is also much larger for *B. terrestris* (Figure 5). The genes *Creld* (LOC100650350) and *aCOP* (LOC100648009 and LOC125386243, 2 copies) are within the BtCSD region and highest heterozygosity levels were found just downstream of *Creld* (based on its orientation), similar to the variation found in *O. biroi* and *O. bicornis*. Blasting of peptide sequences of the flanking genes of BtCSD (*THUMPD3* (LOC100650110)/*Creld*/*aCOPI* the only coding elements) detected wider conservation of this locus in a range of Hymenoptera genome assemblies. The region is conserved across all Aculeata, except the non-CSD ant *Cardiocondyla obscurior*, the three tested Vespinae species and one halictid bee (Table S1). This broad conservation is not detected in the more basal infraorder Proctrupomorpha. The synteny of *THUMPD3*/*Creld*/*aCOP* has been lost and the location of the homologs is at least 0.6Mb apart (Supplemental Table S1). The exception is *Bracon brevicornis*, where all three genes are located in a region of 106 Kb.

**Figure 5.**
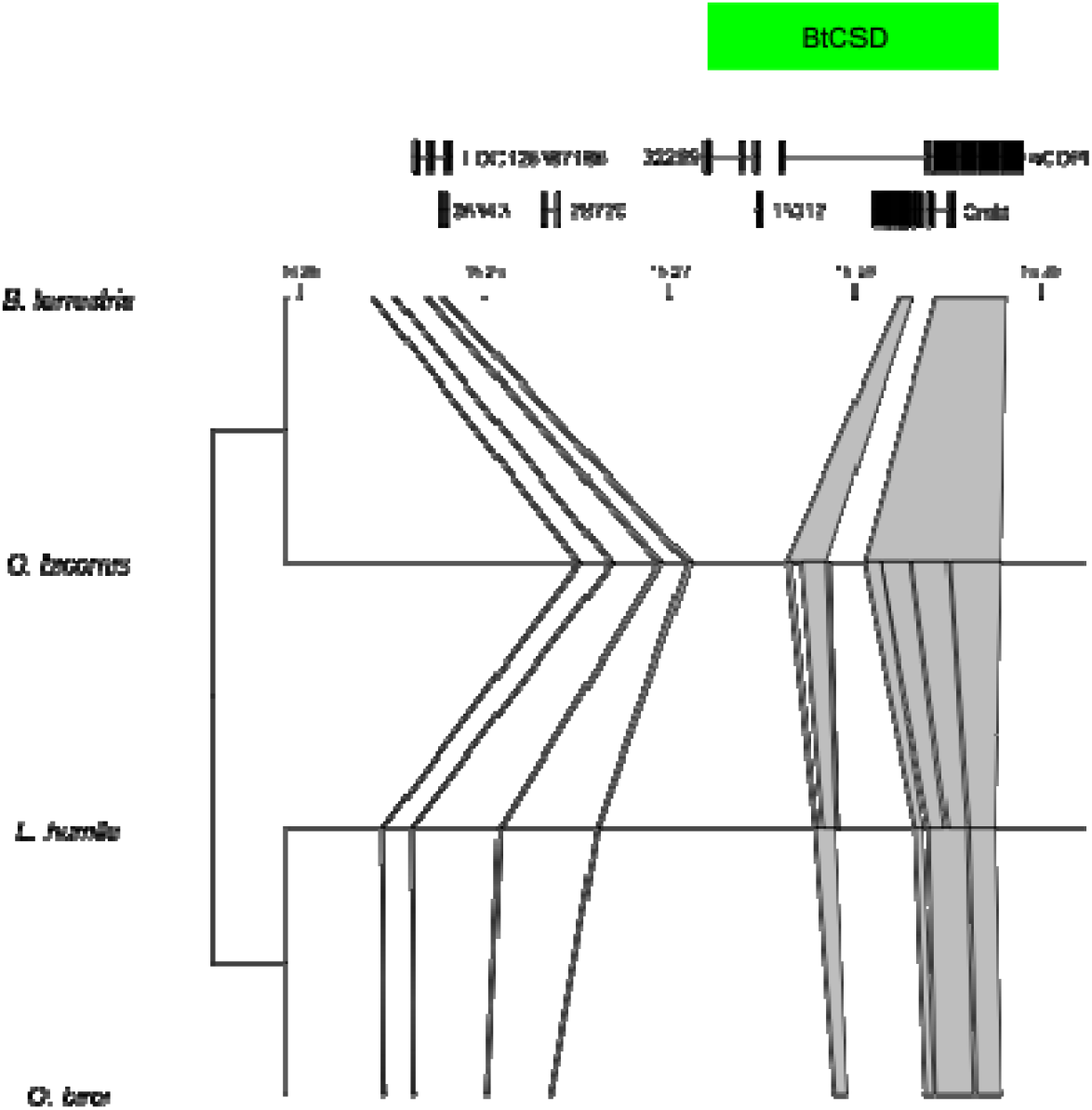
Homology among the CSD loci of *B. terrestris, Osmia bicornis, Linepthema humile* and *Ooceraea biroi*. Lines and blocks indicate places in the locus that are shared among all four species. Green box indicates the positions identified by overlapping low heterozygosity levels at BtCSD (same position as green box figure 2). Numbers on top of the figure indicate positions on Chromosome 2 in Mb.

## Discussion

### The evolutionary implications of an ancestral hymenopteran sex determination mechanism

Despite universally controlling maleness versus femaleness, insect sex determination cascades undergo rapid evolution and diversification (Verhulst et al., 2010; Beukeboom & Perrin, 2014). Con-generics and even con-specifics do not always have the same mechanism. For example, *Musca domestica* houseflies have a base XY system, but different populations have different numbers of male sex-determiner genes on different autosomes (Li et al., 2022). Instructor signals (such as any CSD locus) are particularly variable and fast-evolving (Guerra & Verhulst, 2025). Our study detected conservation between BtCSD, annotated CSD regions of ants (Miyakawa & Mikheyev, 2015; Pan et al., 2024; Lacy et al., 2025) and an orchard bee (Rönneburg et al., 2025), and homologous regions of diverse hymenopteran taxa including other bumblebees (Supplemental Table S1). Furthermore, this *Bombus* CSD candidate shares no homology with the honey bee (*A. mellifera) csd*, a close relative of bumblebees. In *A. mellifera, csd* is a paralog of the conserved insect transducing sex signal *transformer* (called *feminizer* in honey bees), with a hypervariable region corresponding to high allelic diversity (Beye et al., 2003; Beye, 2004). While there are related genes in *B. terrestris* (*transformer* and its paralog *feminizer1*), we and others (Sadd et al., 2015) have ruled these out as CSD candidates because they are not universally heterozygous in diploid females and do not have a hypervariable region, as the region we report here is.

This collectively suggests that BtCSD has deep evolutionary ancestry and represents a conserved Aculeata CSD mechanism (Figure 5). *Apis* CSD, and other possible candidates of *Bracon brevicornis* (Ferguson et al., 2020) and *Lysephebius fabrum* (Matthey-Doret et al., 2019) parasitoid wasps, would then be independent branches of CSD evolution. This reshapes the evolutionary framework of Hymenopteran CSD, as it means inferences from *Apis* as the best-studied system are not broadly applicable. The rapid evolution of new *csd* alleles in the *Apis* hypervariable region corresponds to hundreds of variants worldwide, which is partly due to variation in the number of asparagine/tyrosine repeats (Lechner et al., 2013; Zareba et al., 2017; Seiler & Beye, 2024; Ihle et al., 2025). While this has some superficial similarity to the structural variation of BtCSD, BtCSD’s hypervariable region needs to be assessed for whether it has comparable, or lower de-novo mutation rate and shared allelic diversity across populations. The latter would greatly reduce the potential of genetic rescue in diploid-male producing bottlenecked populations. However, our study’s finding of a distinct phylogenetic origin of BtCSD from *Apis csd* indicates that taxonomic proximity should not be considered predictive of CSD conservation, nor similarity of downstream consequences for population evolution and structure.

### *Factoring in* Bombus *CSD in wild bumblebee conservation*

There are multiple hypotheses as to why bumblebees are suffering more severe losses than other insect and bee groups. One is that many species are ground nesters, and these sites have been greatly reduced due to industrial agriculture (Cameron & Sadd, 2020). Another considers how some stable species such as *B. terrestris* and *B. lucorum* are short-tongued generalist foragers; the loss of longer-tongued species might thus be linked to the specialization to more specific range of flowers, e.g. native species (Goulson & Darvill, 2004). Given its leverage on the detrimental effects of inbreeding in bumblebees, CSD may add to the impacts of these and other factors. Identification of a *Bombus* CSD here paves the way to assessing its role in conservation status, including remedial possibilities (Lecocq et al., 2017).

Since our study suggests likely conservation of *Bombus* CSD, a foremost objective is measuring the number of CSD alleles in threatened species, as its diversity predicts survival prospects. Documentation of high diploid incidence in populations of once-common North American *B. affinis* strongly implicates CSD diversity loss in its decline for example (Mola et al., 2024). CSD diversity measurement will identify populations for which conservation efforts should be prioritized due to low CSD diversity, and donor populations needed for novel CSD allele infusion (accounting for possible outbreeding depression effects; (Gerloff & Schmid-Hempel, 2005). Unfortunately, it may also identify populations that cannot be saved because the minimum level of CSD diversity to sustain a population no longer exists. These designations need to be made carefully, however, as historical context is needed to establish what sustainable levels of CSD diversity are. For example, general inbreeding in *B. veteranus* in the Netherlands, where it is now critically endangered (Slikboer et al., 2023), was indeed confirmed using museum specimens (Maebe et al., 2013). Targeted sequencing, e.g. some marker-level assay for *Bombus* CSD, would give a higher resolution answer on how CSD diversity impacts population health over time. An important observation of our study however is that the colonies inbred for BtCSD were outbred (had high heterozygosity) throughout the rest of genome (Figures 1 and 2). This underscores the importance of knowing CSD loci and measuring it directly, as random genome-wide markers are a poor proxy for inferring diversity for this single gene with outsized impact on population health.

### *Factoring* Bombus *CSD in producing bumblebees*

Currently, commercial bumblebee production of commonly used *B. terrestris* and *B. impatiens* seems to be stable (Osterman et al., 2021). The exact costs of unusable diploid male-producing colonies are not disclosed by commercial breeders. However, by destroying these colonies, they are performing a human-mediated form of CSD diploid male vortex mitigation. In future, commercial breeders may electively evaluate the CSD-related aspects of population health in their stocks by measuring and monitoring the amount of CSD allelic diversity. Even with diploid-male colony destruction, if the initial pool of CSD alleles is insufficient and crosses not sufficiently variable, an isolated captive breeding stock may have a limited “lifespan”.

Critically, knowledge of this *Bombus* CSD candidate will be essential to developing new local bumblebee strains or species for large-scale agricultural pollination. The rapidly growing interest in this industry is motivated by a desire to reduce reliance on *B. terrestris* imports, which at times has led to resource competition with native bees including other bumblebees (Ishii et al., 2008; Williams & Osborne, 2009; Goka, 2010); disease spillover of pathogens and parasites (Smith-Ramírez et al., 2023); and introgression into native populations (Kardum Hjort et al., 2022; Franchini et al., 2023). The last endangers native genetic diversity and its linkage to local adaptation; it can also produce sterile hybrids (Kondo et al., 2009) However, poor management of CSD diversity may prevent successful development of new commercial populations. This candidate may be used to design new breeding protocols that ensure sufficient CSD alleles in a starting population; crosses that maintains this diversity and prevents any costly diploid male production; and identifies sources of distinct alleles for genetic rescue if needed (e.g. different wild populations).

### *Functional investigation of* Bt*CSD*

A precursor to any of the above fundamental and applied avenues for this Bombus CSD candidate is more mechanistic knowledge. Importantly, *BtCSD* still requires functional validation. As CSD is a feminization signal, preventing its expression in heterozygous embryos through RNA interference would divert diploid female development to male development (e.g. as with other hymenopteran sex determiners: (Zou et al., 2020; Pan et al., 2024). The exact target(s) in the large 2Kb region are still to be determined. While a *Bombus* CSD candidate was previously suggested using a *B. terrestris* linkage map, it was derived from randomly distributed RAPD markers (Wilfert et al., 2006; Stolle et al., 2011) and could not be mapped back here to narrow a region of interest. However, a hint may come from the ant *L. humile*, as targeting the non-coding region in this ant resulted in only a weak knockdown effect (expression of male splice *transformer* in ~10% of treated heterozygous eggs) (Pan et al., 2024). This long non-coding RNA does not show any homology to the BtCSD and CSD locus of *Osmia bicornis*. This suggests that lncRNA candidates are linked to CSD loci rather than being CSD outright. As such, the most promising candidate targets could be the downstream coding genes *Creld* and the two copies of a*COP* in the hypervariable region. *Creld* is positioned with its C-terminal region extending into the structurally variable region (Figure 3). A proper functional testthat elucidates the genetic element for functional sex determination in BtCSD (i.e. a knockdown-or-knockout of a candidate that results in heterozygous male development) will allow for a better comparison between the bumblebee and the ant CSD locus.

Once the functional gene(s) have been identified, a major aspect of the functionality of CSD is the amount of sequence difference needed to qualify different alleles. For example, the amount of difference needed between alleles for heterozygosity (heterotrimer formation) versus homozygosity (homotrimer formation) is still debated even within the well-studied and high-CSD diversity *Apis* system (Otte et al., 2023). There is a complex model requiring both differences of length and a threshold number of amino acid mismatches in the hypervariable region (Lechner et al., 2013); and simpler models arguing that a minimum of 3-5 amino acid differences (Beye et al., 2013), or even one (Mroczek et al., 2022), is sufficient. Testing this for bumblebees would require controlled crosses with alleles falling across a gradient of relatedness. This in turn needs sequencing of CSD candidate to cross individuals with the right allelic combinations.

As a synthesis, identification of *Bombus* CSD is the launching point for many branches of bumblebee research. This ranges from to better definition of the extinction risk of CSD in *Bombus* population dynamics and remedial possibilities; to improved breeding of existing and to-be-founded commercial populations; to further delineation of the mechanism itself.

## Supporting information

Supplemental Table S1 References

## Acknowledgements

This work was funded by the Dutch Research Council (NWO) with VI.Veni.232.066 awarded to Kelley Leung. We thank our commercial partner for providing materials for sequencing. The authors declare no conflict of interest.

## Author Contributions

Joost van den Heuvel, Kelley Leung and Bart A. Pannebakker conceived this study. Elzemiek Geuverink, Peter Šima, Thomas V. M. Groot and Roland Kreskóci advised on its design. Peter Šima, and Roland Kreskóci provided specimens. Frank Becker and Joost van den Heuvel collected data. Joost van den Heuvel and Elzemiek Geuverink analyzed data. Kelley Leung secured funding. Kelley Leung and Joost van den Heuvel wrote the initial draft of the manuscript. All authors discussed results and contributed to the writing of other versions.

## Data availability statement

All raw data have been uploaded to the NCBI Short Read Archive in Bioproject PRJNA1296802.

## Supplemental figures

**Figure S1.**
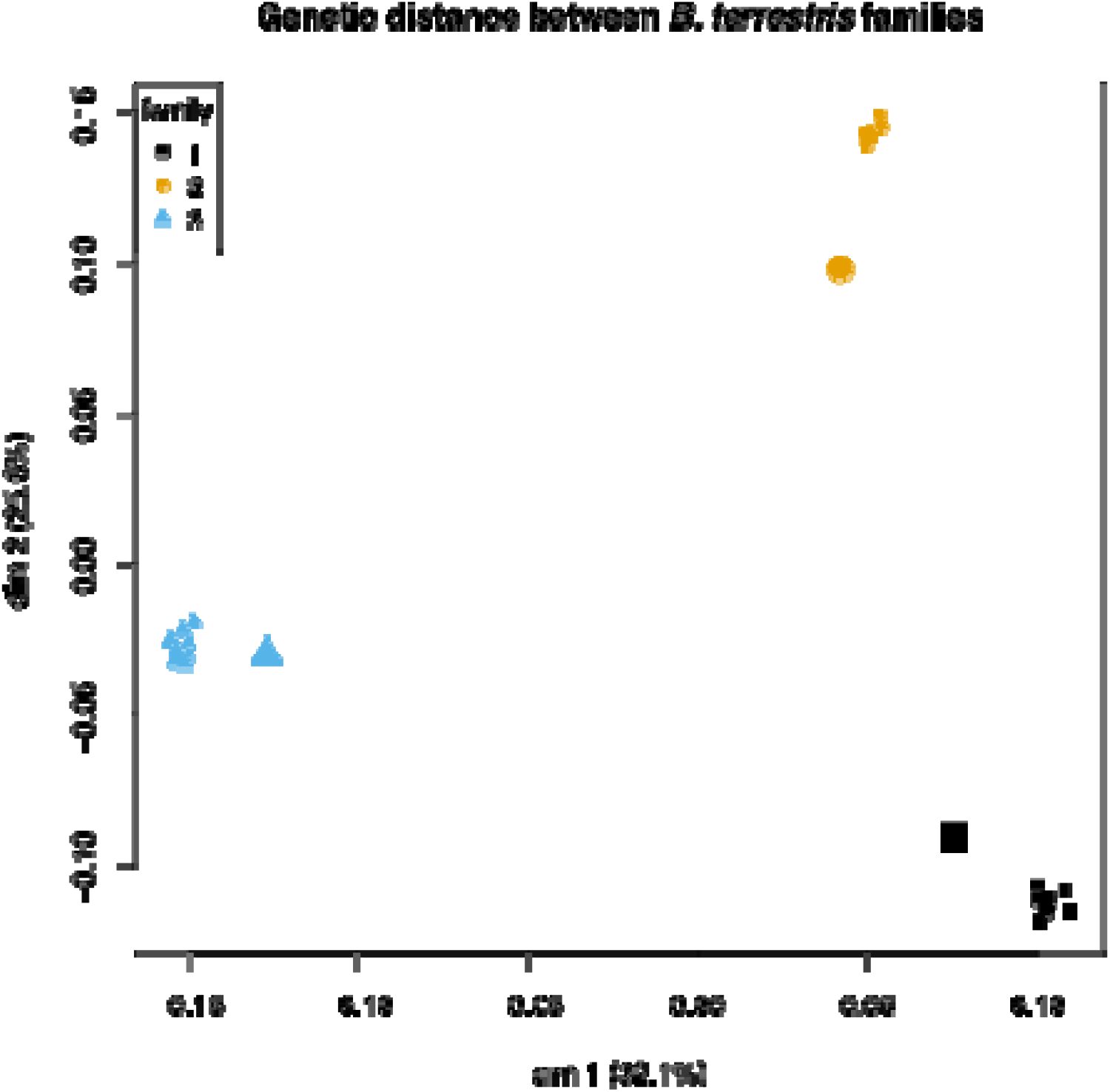
Genetic distance between sampled *B. terrestris* colonies. Multidimensional scaling of genetic distance among the 33 samples, including 1 queen and 10 diploid males per family. Dimension 1 explains 32.1% of the total variation, while 25.6% is explained by dimension 2. Different colonies are indicated by varying symbols and color. The larger symbols indicate queens. This plot is indicative of all family members being genetically relatively close to each other.

**Figure S2.**
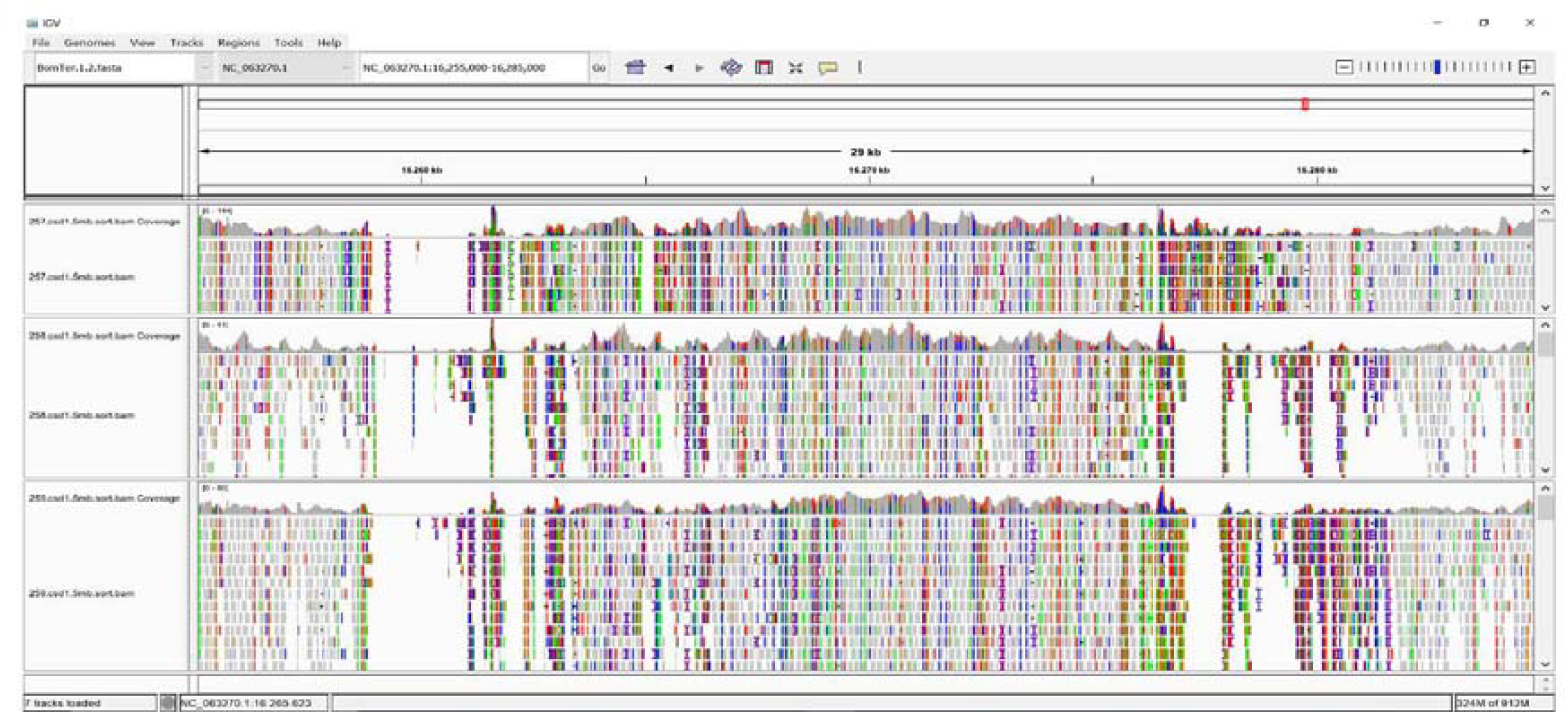
Integrative genome viewer output for the CSD locus on Chromosome 2. From top to bottom alignments of the queens of Family 1, 2 and 3 are shown. Alignment is lacking around 16,260 Kb, but also partly around 16,265 for Queen 1 in the upstream area. In the downstream area coverage is lacking between 16,275 and 16,280 Kb. The colors in each upper panel (coverage) panels indicate variation compared to the reference genome. Very clearly, a lot of genomic variation is inferred around the regions with missing coverage.

**Figure S3.**
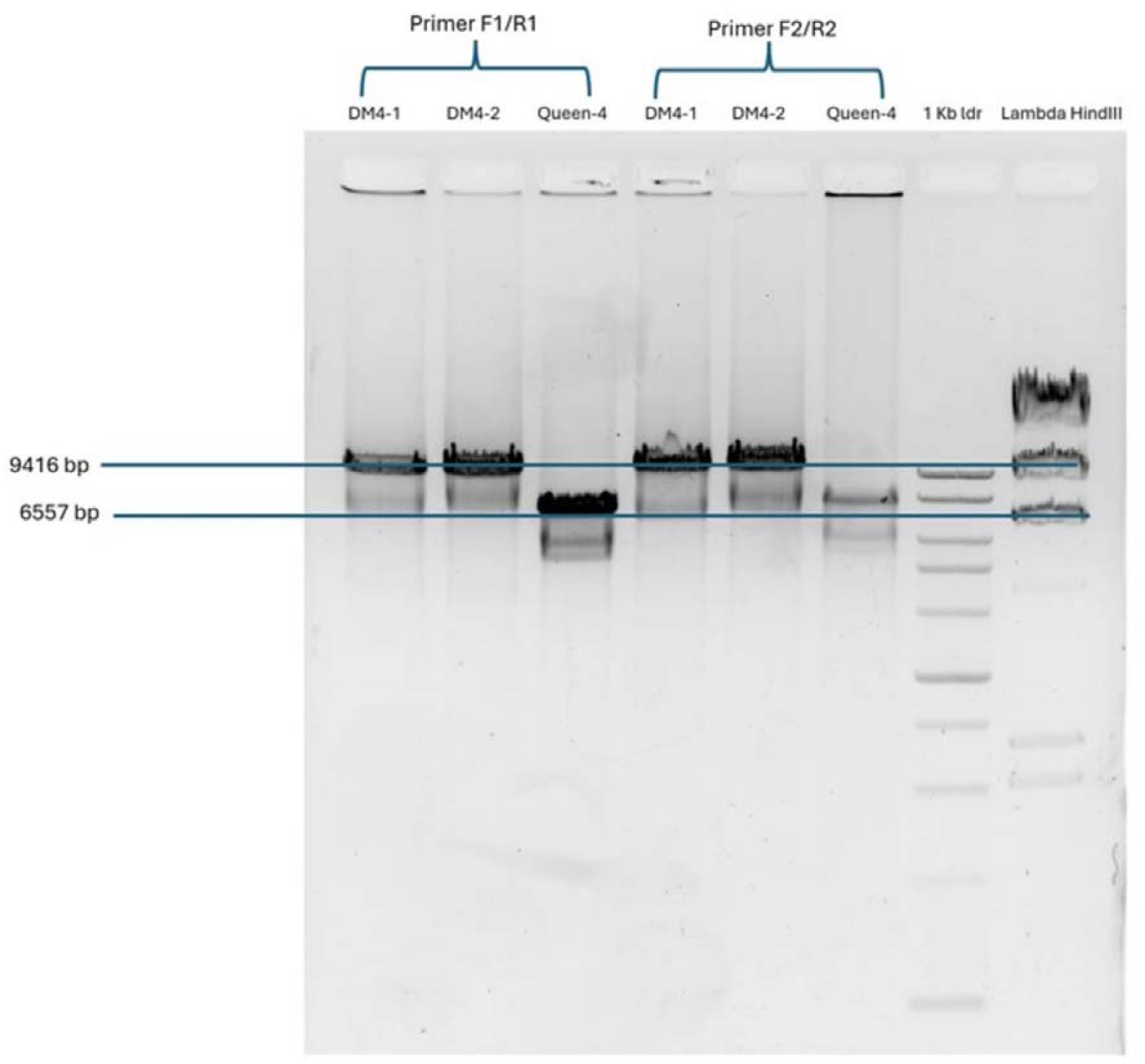
Gel electrophoresis result from long distance PCR on the queen and two diploid males from Family 4 produced one long fragment in two diploid males (dm4-1 and dm4-2) and one shorter fragment in the queen (Q4). The lengths of two separate primer pair amplified seemingly the same region.

Table S1. Detection of THUMPD3/Creld/acOPI genomic region across Hymenoptera. Species were selected based on availability of high-quality genome, (if known) presence/absence of CSD mechanism and phylogenetic position. (Not included here because of size; see attached excel file)

