## Supplemental File 3 for "*Bombus terrestris* Complementary Sex Determiner (BtCSD) is identified as a conserved hymenopteran sex determination region: evolution, breeding, and conservation implications": supplemental file 3 kmers based synthblock determination.R

```
files<- list.files()
files<-files[which(substring(files,nchar(files)-5,nchar(files))==".fasta")]

files

pops<-unlist(strsplit(files, ".fasta"))

library(seqinr)

### first we need to determine which kmers are usefull to group 
### sequences

U<-c()
for (I in 1:length(pops))
{

fasta<- read.fasta(paste(pops[I],".fasta", sep=""))
kmers<-c()
L1<- length(fasta)

for (i in 1:L1)
{
if (length(fasta[[i]])>1000){
for (j in 1:(length(fasta[[i]])-15))
{
seq<- fasta[[i]][j:(j+14)]
rseq<- rev(comp(seq))
kmers<- rbind(kmers,sort(c(getSequence(seq, as.string=TRUE)[[1]][1],
getSequence(rseq, as.string=TRUE)[[1]][1]))[1])
}}}
U<- c(U, kmers)
plot(sort(as.numeric(table(U))),main=I)

}


write.table(names(which(table(U)>2 & table(U)<22)),
file="common kmers.txt", col.names=FALSE, row.names=FALSE, quote=FALSE)


U<-names(which(table(U)>2 & table(U)<22))

Score<-rep(0, length(U))

for (i in 1:length(U))
{

if(grepl("w",U[i]) | grepl("n",U[i]) | grepl("x",U[i]) | grepl("NA",U[i])){
Score[i]<- 1}
}

U<- U[which(Score==0)]

write.table(U,
file="common kmers.txt", col.names=FALSE, row.names=FALSE, quote=FALSE)

### now we save the kmers and grouping....


for (I in 1:length(pops))
{

fasta<- read.fasta(paste(pops[I],".fasta",sep=""))
kmers<-c()
L1<- length(fasta)

for (i in 1:L1)
{
if (length(fasta[[i]])>500){
for (j in 1:(length(fasta[[i]])-15))
{
seq<- fasta[[i]][j:(j+14)]
rseq<- rev(comp(seq))
kmers<- rbind(kmers,c(sort(c(getSequence(seq, as.string=TRUE)[[1]][1],
getSequence(rseq, as.string=TRUE)[[1]][1]))[1],i,j))
}}}

kmers<-kmers[which(kmers[,1] %in% U),]

GR<-1
for (i in 2:length(kmers[,1]))
{
if (kmers[i-1,2]==kmers[i,2] & (as.numeric(kmers[i,3])-as.numeric(kmers[i-1,3]))<20)
{
GR<-c(GR,GR[i-1])
}else {GR<- c(GR,GR[i-1]+1)}
}

write.table(file=paste(pops[I],".kmers.txt",sep=""), cbind(kmers,GR),
col.names=FALSE, row.names=FALSE, quote=FALSE, sep='\t')
plot(GR,main=I)
}


comdat<-read.table("common kmers.txt")$V1

for (i in 1:length(pops))
{
kmers<- read.table(paste(pops[i],".kmers.txt", sep=""), header=F)
L<-rep(0, length(comdat))
for (j in 1:length(kmers[,1]))
{
W<- which(comdat==kmers[j,1])

if (length(W)==1)
{
L[W]<- kmers[j,4]
}
}
plot(L, main=i)

write.table(L, file=paste(pops[i],"knum.txt", sep=""),
col.names=FALSE, row.names=FALSE, quote=FALSE, sep='\t')

}


totdat<- as.character(read.table(paste(pops[1],"knum.txt", sep=""),header=F)$V1)

for (i in 2:length(pops))
{
dat<-read.table(paste(pops[i],"knum.txt", sep=""),header=F)$V1
totdat<-paste(totdat,dat)
}

unique(totdat)

Udat<- unique(totdat)

L<-c()
for (i in 1:length(Udat))
{
L<- c(L,sum(totdat==Udat[i]))
}

plot(L)

GRsize<-c()
for (i in 1:length(Udat))
{
GRsize<-c(GRsize,length(pops) - sum(strsplit(Udat[i]," ")[[1]]==0))
}

plot(GRsize ~ L)


CANDI<-Udat[which(L>400)]

CANDI


MAT<- matrix(0L, ncol=length(pops), nrow=length(pops))

for (i in 1:(length(pops)-1)){
for (j in (i+1):length(pops)){
dat1<-read.table(paste(pops[i],"knum.txt", sep=""),header=F)$V1
dat2<-read.table(paste(pops[j],"knum.txt", sep=""),header=F)$V1

dat1[which(dat1>0)]<-1
dat2[which(dat2>0)]<-1

MAT[i,j]<-1- (sum(dat1==1 & dat2==1) /min(c( sum(dat1),sum(dat2))))
MAT[j,i]<- MAT[i,j]
}
}


colnames(MAT)<- pops
rownames(MAT)<- colnames(MAT)

#MAT<-MAT[c(1:7,20,21),c(1:18,20,21)]

MAT<-MAT[c(1:7,10:16,18:21),c(1:7,10:16,18:21)]


which(colnames(MAT)

library(ape)

pdf("genetic kmer cluster.pdf",useDingbats=FALSE)
plot(as.phylo(hclust(as.dist(MAT))),cex=0.6)
dev.off()
YORDER<-hclust(as.dist(MAT))$order

WHATseq[which(WHATseq[,2]==45),]

################# first we determine which scaffolds should be where


### correcting YORDER to remove previous 18
#YORDER<-c(YORDER,18)
#YORDER[12]<- YORDER[12]+1
##YORDER[14]<- YORDER[14]+1
#YORDER[20]<- YORDER[20]+1


SIZE<-400

WHATseq<-c()
for (N in 1:length(pops)){
fasta<- read.fasta(paste(pops[N],".fasta",sep=""))
for (K in 1:length(which(L>SIZE))){
kmers<- read.table(paste(pops[N],".kmers.txt", sep=""), header=F)
if (as.numeric(strsplit(Udat[which(L>SIZE)[K]]," ")[[1]])[N]>0)
{
WWW<-which(kmers[,1] %in% comdat[which(totdat==Udat[which(L>SIZE)[K]])])
P<-as.numeric(names(which(table(kmers[WWW,4])==max(table(kmers[WWW,4])))))
M<-kmers[which(kmers[,1] %in% comdat[which(totdat==Udat[which(L>SIZE)[K]])] & kmers[,4]==P),3]
S<- table(kmers[which(kmers[,1] %in% comdat[which(totdat==Udat[which(L>SIZE)[K]])] & kmers[,4]==P),2])
S<-as.numeric(names(which(S==max(S))))
WHATseq<- rbind(WHATseq,c(N,K,S,length(fasta[[S]]), min(M),max(M)))
}}}


WHATseq<- WHATseq[which(WHATseq[,1] %in% c(1:7,10:16,18:21)),]

WHATseq[which(WHATseq[,1]==10),1]<-8
WHATseq[which(WHATseq[,1]==11),1]<-9
WHATseq[which(WHATseq[,1]==12),1]<-10
WHATseq[which(WHATseq[,1]==13),1]<-11
WHATseq[which(WHATseq[,1]==14),1]<-12
WHATseq[which(WHATseq[,1]==15),1]<-13
WHATseq[which(WHATseq[,1]==16),1]<-14
WHATseq[which(WHATseq[,1]==18),1]<-15
WHATseq[which(WHATseq[,1]==19),1]<-16
WHATseq[which(WHATseq[,1]==20),1]<-17
WHATseq[which(WHATseq[,1]==21),1]<-18

write.table(WHATseq, 
file="WHATseq.txt", col.names=c("ind","syntblock","scaf","length","start","end"),
row.names=FALSE, quote=FALSE, sep='\t')

WHATseq<- read.table("WHATseq.txt", header=T)


TAB<-tapply(WHATseq[,5],WHATseq[,2],mean)


COMTAB<-which(table(WHATseq[,2])==18)


MINGROUP<-as.numeric(names(which(TAB[which(names(TAB) %in% COMTAB)]==min(TAB[which(names(TAB) %in% COMTAB)]))))
LEVELMING<- as.numeric(TAB[which(names(TAB)==MINGROUP)])


plot(-10000000,-100000,xlim=c(10000,55000),
ylim=c(1,length(pops)), xlab="relative position", ylab="individual",main="structral variation")


library(RColorBrewer)
color = grDevices::colors()[grep('gr(a|e)y', grDevices::colors(), invert = T)]
COLS<- sample(color,max(WHATseq[,2]))
length(COLS)

write.table(COLS, file="synthcolors.txt",col.names=FALSE, row.names=FALSE,
sep='\t', quote=FALSE) 

SYNTHsize<- WHATseq[,6]-WHATseq[,5]

plot(SYNTHsize)

WHATseq<- WHATseq[which(SYNTHsize<5000),]

table(WHATseq[,2])

### we monitor the start of 2, and then the next round we can correct X values

WHATseq[which(WHATseq[,1]==9),]
WHATseq[which(WHATseq[,1]==8),]
WHATseq[which(WHATseq[,1]==17),]

which(YORDER==18)

LET<- c(letters,toupper(letters[1]))


WHATseq[,2]

pdf("syntheny blocks.pdf", useDingbats=FALSE, width=8, height=8)

plot(-10000000,-100000,xlim=c(0,55000),
ylim=c(1,18), xlab="relative position", ylab="",main="structural variation CSD locus",
yaxt="n")

for (i in 1:length(WHATseq[,2]))
{
Y<- YORDER[WHATseq[i,1]]
Xstart<-WHATseq[i,5]
Xend<- WHATseq[i,6]

if (Y==17){
Xstart<- Xstart
Xend<- Xend
}

if (Y==18){
Xstart<- Xstart
Xend<- Xend
}


WW<- which(WHATseq[,1]==WHATseq[i,1] & WHATseq[,2]==MINGROUP)
if (length(WW)>0){
Xstart<- Xstart - WHATseq[WW,5]+LEVELMING
Xend<- Xend - WHATseq[WW,5]+LEVELMING
}
rect(Xstart,Y-0.4, Xend,Y+0.4, col=COLS[WHATseq[i,2]],border=FALSE)
COLO<- "black"
if (WHATseq[i,2]==7 | WHATseq[i,2]==14 | WHATseq[i,2]==25){COLO<- "white"}
text((Xend+Xstart)/2,Y,LET[WHATseq[i,2]],cex=0.8,col=COLO)
}

axis(2, at=c(-100,100),col="white")


dev.off()
```
