## Supplemental Table S1 References for "*Bombus terrestris* Complementary Sex Determiner (BtCSD) is identified as a conserved hymenopteran sex determination region: evolution, breeding, and conservation implications"

**References Supplementary Table**

(Hasselmann *et al.*, 2008) 10.1093/molbev/msn011

(Beye *et al.*, 2003) https://doi.org/10.1016/s0092-8674(03)00606-8

(Pamilo *et al.*, 1994) https://doi.org/10.1080/08927014.1994.9522996

(Schrempf *et al.*, 2006) 10.1038/sj.hdy.6800846

(Ross & Fletcher, 1985) https://doi.org/10.2307/2408688

(Foster *et al.*, 2000) https://doi.org/10.1046/j.1365-294x.2000.00920.x

(Speicher & Speicher, 1940) https://doi.org/10.1086/280904

(Gu & Dorn, 2003) https://doi.org/10.1006/anbe.2003.2185

(Steiner & Teig, 1989) https://www.cabidigitallibrary.org/doi/full/10.5555/19891132952

(Butcher *et al.*, 2000) https://doi.org/10.1046/j.1420-9101.2000.00203.x

(Beukeboom, 2001) https://doi.org/10.1163/156854201X00017

Beukeboom, L.W. (2001) Single-locus complementary sex determination in the ichneumonid Venturia Canescens (Gravenhorst) (Hymenoptera). *Netherlands Journal of Zoology*, **51**, 1–15.

Beye, M., Hasselmann, M., Fondrk, M.K., Page, R.E. & Omholt, S.W. (2003) The gene csd is the primary signal for sexual development in the honeybee and encodes an SR-type protein. *Cell*, **114**, 419–429.

Butcher, R.D.J., Whitfield, W.G.F. & Hubbard, S.F. (2000) Complementary sex determination in the genus Diadegma (Hymenoptera: Ichneumonidae). *Journal of Evolutionary Biology*, **13**, 593–606.

Foster, K.R., Ratnieks, F.L.W. & Raybould, A.F. (2000) Do hornets have zombie workers? *Molecular Ecology*, **9**, 735–742.

Gu, H. & Dorn, S. (2003) Mating system and sex allocation in the gregarious parasitoid *Cotesia glomerata*. *Animal Behaviour*, **66**, 259–264.

Hasselmann, M., Vekemans, X., Pflugfelder, J., Koeniger, N., Koeniger, G., Tingek, S., *et al.* (2008) Evidence for convergent nucleotide evolution and high allelic turnover rates at the complementary sex determiner gene of Western and Asian honeybees. *Molecular Biology and Evolution*, **25**, 696–708.

Pamilo, P., Sundström, L., Fortelius, W. & Rosengren, R. (1994) Diploid males and colony-level selection in Formica ants. *Ethology Ecology & Evolution*, **6**, 221–235.

Ross, K.G. & Fletcher, D.J.C. (1985) Genetic Origin of Male Diploidy in the Fire Ant, Solenopsis invicta (Hymenoptera: Formicidae), and its Evolutionary Significance. *Evolution*, **39**, 888.

Schrempf, A., Aron, S. & Heinze, J. (2006) Sex determination and inbreeding depression in an ant with regular sib-mating. *Heredity*, **97**, 75–80.

Speicher, B.R. & Speicher, K.G. (1940) The Occurrence of Diploid Males in Habrobracon brevicornis. *The American Naturalist*, **74**, 379–382.

Steiner, W.W.M. & Teig, D.A. (1989) Microplitis croceipes (Cresson): Genetic characterization and developing insecticide resistant biotypes. *Southwestern Entomologist*, **12**, 71–87.
